# DENetwork: Unveiling Regulatory and Signaling Networks Behind Differentially-Expressed Genes

**DOI:** 10.1101/2023.06.27.546719

**Authors:** Ting-Yi Su, Quazi S. Islam, Steven K. Huang, Carolyn J. Baglole, Jun Ding

**Author notes:** To whom correspondence should be addressed. Email: CJB or JD.

## Abstract

Differential gene expression analysis from RNA-sequencing (RNA-seq) data offers crucial insights into biological differences between sample groups. However, the conventional focus on differentially-expressed (DE) genes often omits non-DE regulators, which are an integral part of such differences. Moreover, DE genes frequently serve as passive indicators of transcriptomic variations rather than active influencers, limiting their utility as intervention targets. To address these shortcomings, we have developed *DENetwork*. This innovative approach deciphers the intricate regulatory and signaling networks driving transcriptomic variations between conditions with distinct phenotypes. Unique in its integration of both DE and critical non-DE genes in a graphical model, *DENetwork* enhances the capabilities of traditional differential gene analysis tools, such as *DESeq2*. Our application of *DENetwork* to an array of simulated and real datasets showcases its potential to encapsulate biological differences, as demonstrated by the relevance and statistical significance of enriched gene functional terms. *DENetwork* offers a robust platform for systematically characterizing the biological mechanisms that underpin phenotypic differences, thereby augmenting our understanding of biological variations and facilitating the formulation of effective intervention strategies.

## INTRODUCTION

RNA sequencing (RNA-seq) has emerged as a prominent genomics technology that is extensively used to profile cellular states in high-throughput fashion. A key task in the analysis of RNA-seq data involves the systematic investigation (i.e., on a whole transcriptome scale) of biological differences between samples under various conditions, often linked to distinct phenotypes. To illustrate, RNA-seq measurements are widely employed to discern transcriptomic differences between samples from healthy controls and patients with diverse diseases (1, 2, 3). The understanding of such biological differences is instrumental in identifying biomarkers for accurately diagnosing complex diseases (4, 5), and is a fundamental prerequisite for the development of therapeutic drugs.

Identifying differentially-expressed (DE) genes between conditions has been the most prevalent strategy for pinpointing biomarkers. This technique has demonstrated considerable success across a broad spectrum of biological problems, including complex diseases (6, 7, 8, 9, 10, 11). Over the past decade, a myriad of methods for discerning DE genes between samples under different conditions have been developed. Techniques such as DESeq2 (12) and edgeR (13) are the most widely adopted for detecting DE genes from RNA-seq data, and have undeniably enhanced our understanding of the mechanisms driving countless biological processes and diseases (4, 5, 14).

Notably, differential gene analysis methods are also adaptive and capable of accommodating new data types and research questions. Extensions and variants of these methods have been applied to detect DE genes from the rapidly growing field of high-resolution single-cell RNA-seq data. This newer form of data allows researchers to profile all the cells in a sample, and differential gene analysis methods have demonstrated success in various application scenarios (15, 16, 17, 18).

However, while differential gene analysis methods have catalyzed advancements in various biological fields, they have their limitations. They primarily showcase partial differences mirrored by the change in gene expression between conditions. More often than not, DE genes are the effects rather than the causes of the biological disparities between conditions. Hidden beneath surface-level changes are the gene regulatory and signaling networks that orchestrate the observed gene expression changes. One critical aspect of these networks involves cellular receptors, specialized proteins located on the cell surface or within the cell, which facilitate cell communication and environmental sensing. They bind to specific signaling molecules, or ligands, such as proteins, peptides, hormones, or small molecules from neighboring cells or the extracellular environment (19, 20). Ligand binding triggers a cascade of intracellular events, often leading to changes in gene expression, followed by post-transcriptional events regulated by RNA-binding proteins. Such events induce dynamic changes in protein expression that are vital for cellular functions. This process, known as signal transduction, enables cells to respond and adapt to changes in their surroundings and facilitates communication between cells. If the objective is to identify potential intervention targets to manipulate biological differences (e.g., modifying disease samples to make them resemble healthy ones), then the transcription factors (TFs), signaling proteins, and receptors involved in the signal transduction process become increasingly important.

Unfortunately, to the best of our knowledge, there is a dearth of tools capable of comprehensively delineating biological discrepancies between conditions, particularly those that extend beyond merely DE genes and offer additional perspectives, such as signal transduction. To bridge this gap, we introduce a unified framework, DENetwork. This system aims not only to identify DE genes but also to reconstruct the underlying regulatory and signal networks driving differential gene expression. To augment the utility of the reconstructed networks for identifying potential intervention targets, we also devised and implemented an in-silico knockout strategy. This method seeks to score and rank all network nodes, which include DE genes, TFs, signaling proteins, and receptors. The nodes that rank highly are indicative of critical genes accounting for the biological distinctions between conditions. In particular, the top-ranked receptors and TFs may serve as potential candidates for future interventions, perhaps paving the way for advancements in therapeutics.

## MATERIALS AND METHODS

### Data and data pre-processing

In this study, we leveraged three publicly available bulk RNA-seq datasets, each consisting of distinct biological conditions, to evaluate the performance of our method. These datasets encompass gene expression measurements in response to Influenza A virus (IAV) infection (GSE192528), SARS-CoV-2 infection (21), and the 12-lipoxygenase and 15-lipoxygenase (Alox15-/-) genetic deficiencies in mouse macrophages (22). Moreover, our proprietary bulk RNA-seq dataset that investigates the role of HuR (human antigen R) in human lung fibroblasts was also subjected to this method (23, 24). For our analysis, the IAV dataset, which originally comprised RNA-seq data of human lung tissue following infections from IAV, Pseudomonas aeruginosa, and Mycobacterium bovis (BCG), was narrowed down to include only five IAV-infected samples and five uninfected controls. Given the imbalance (36 positive samples and 5 negative samples) in the SARS-CoV-2 dataset, we employed UMAP to perform dimensionality reduction and segregate the samples. This resulted in a more balanced subset of 12 positive and four negative samples (Supplementary figure S1). The macrophage dataset comprised three wild-type samples and four samples with Alox15-/-genetic deficiency. And the HuR dataset consisted of three normal human fibroblast samples and three HuR knockdown samples.

The raw gene expression count matrix for each dataset was analyzed using DESeq2 (12), facilitating the identification of upregulated and downregulated DE genes across different conditions. The resultant DESeq2 output was further filtered based on established thresholds: baseMean *>* 50, unadjusted or False Discovery Rate (FDR) adjusted *p*-value *<* 0.05, log2fc *>* 0.6 (for upregulated genes), and log2fc *<* -0.6 (for downregulated genes). The cutoff for DE genes can be adjusted to fit different application scenarios. For example, for the HuR dataset, the baseMean threshold was changed to *>* 500 and the log2fc thresholds were changed to *>* 1.5 and *<* -1.5 for upregulated and downregulated genes, respectively, to obtain a more confident list of DE genes.

Concurrently, we curated receptors specific to each dataset: 200 IAV receptors from the mt-STEM study (25), 332 SARS-CoV-2 receptors derived via our COVID2GeneList method (26), and 158 macrophage receptors obtained from all expressing mouse receptors in the macrophage dataset in CellTalkDB (27). Acknowledging the significance of sing dataset-specific receptors, we sourced all relevant receptors from public receptor databases such as CellTalkDB (27) in cases where specific information was unavailable. Any receptors not expressed within a given dataset were excluded from downstream analyses. This approach was employed for the HuR dataset, where a specific list of receptors was not readily accessible.

### Building an initial fully connected network

In this study, we orchestrated a fully connected network comprising gene nodes, interconnected by edges symbolizing either protein-protein (pp) or protein-DNA (pd) interactions. To reflect disparities between biological conditions, the absolute value of log2 fold changes of the genes were denoted as node attributes. Both pp and pd interaction scores, ranging between 0 and 1, were assigned as edge weights. When both interaction data types were available for a gene pair, the mean value served as the edge weight.

Our network designated receptors as source nodes, given their role in initiating signal transduction, while genes displaying differential expression between conditions constituted the target nodes. Any DE gene that is also a receptor was designated as a source node. In a bid to facilitate users’ exploration of gene regulatory networks underlying gene expression differences, we also included TFs as potential target nodes (Supplementary Figures S5 and S6). We leveraged the TF-gene interaction data for common species provided in the iDREM software (28) and used a binomial test to detect TFs with significant DE target genes.

For every candidate TF, we enumerated all target genes (*n*) and pinpointed DE genes (*x*). The background probability of DE genes, *P*_*bg*_, was calculated as the ratio of DE genes to all genes. Subsequently, we computed the TF enrichment p-value as *p*-value= 1 − *pbinom*(*x −* 1,*n*,*P*_*bg*_). Post FDR or Bonferroni correction, all TFs with an adjusted enrichment p-value *<* 0.05 were enlisted as target TF nodes. The NetworkX Python package (29) served as a fundamental tool for the construction of our network.

### Finding significant shortest paths from source to target

In this step, our goal was to discern the shortest paths *X* between pairs of source (*S*) and target (*T*) nodes within our gene network. A limiting factor was implemented, wherein the number of intermediate nodes along each path must not surpass a predetermined maximum, labeled as *K*, as shown in Equation **1**:

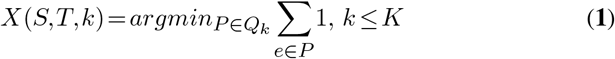

In this equation, *Q*_*k*_ represents all potential paths *P* encompassing *k* intermediate nodes. All edges were assumed to have a weight of 1.

Once these shortest paths were isolated, we attributed a score *s* to each path *P* . This scoring is mathematically depicted in Equation **2**:

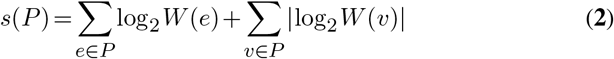

In this equation, *W* (*e*) denotes the PPI score of edge *e*, and *W* (*v*) marks the fold change of node *v*. To simplify calculations and preclude scores from approaching zero, we applied log2 transformations to the scores.

We postulated that these path scores would follow a Gaussian distribution, a hypothesis corroborated by the distribution plot found in Supplementary Figure S2. Under this assumption, we computed a p-value *p* for each path score using Equation **3**:

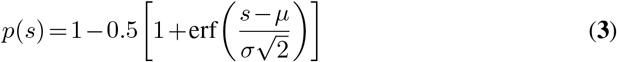

In this equation, *s* corresponds to the path score, while *µ* and *σ* denote the mean and standard deviation of the Gaussian path score distribution, respectively. Any path with a p-value below 0.02 was considered statistically significant and retained for further exploration.

In the final phase of this process, we focused on reducing redundancy and promoting network sparsity. To achieve this, for each pair of source and target nodes in the network, we selected the top *N* paths — also referred to as the most probable paths — with the maximal scores *M*_*p*_ based on Equation **4**:

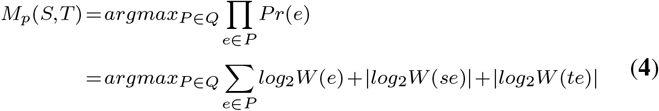

In this equation, *M*_*p*_(*S*,*T*) represents the set of most probable paths for a source and target pair. The probability of a path’s occurrence is determined as the product of the probabilities of the edges in the path. The probability of an edge (*e*) is defined as the product of its PPI score *W* (*e*), the fold change of its starting node *W* (*se*), and the fold change of its ending node *W* (*te*). These products are converted to summations after applying log2 transformations.

We empirically set the hyperparameters *N* and *K* to 5. However, users can adjust these values to better suit their datasets. It is important to note that larger values for *K* and *N* typically allow for a more accurate reconstruction of the signaling network, but also result in an exponential increase in search space, which could lead to a considerable rise in computational complexity.

### Searching for a local-optimal network

In our pursuit of a local-optimal network, we adopted a greedy strategy that involved the systematic elimination of paths, starting from those with the lowest path scores (determined by equation **2**). Following the removal of each path, we generated a new graph, *Q*, and calculated a corresponding score, *S*(*Q*), as illustrated in equation **5**:

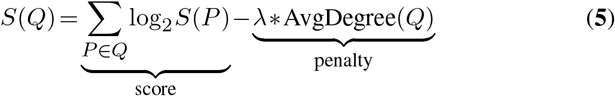

Equation **5** encompasses two components: the score term, which corresponds to the aggregate score of *Q* (sum of the log2-transformed scores of all paths persisting in *Q*), and a penalty term. This penalty term accounts for network complexity and is proportional to the average degree of *Q*. By default, the regularization coefficient *λ* is set to the ratio of *mean*(score term) to *mean*(AvgDegree(*Q*)). We procedurally removed *I* paths with the lowest scores, where *I* was typically set to 50. Following each removal, we graphed *S*(*Q*) against the quantity of removed paths (*i ≤ I*). The overall network score *S*(*Q*) had a tendency of decreasing as more paths were discarded, paralleling a reduction in network complexity. We hence generated a plot between *S*(*Q*) and *i*, and identified the graph *Q* corresponding to the ‘elbow’ or ‘knee’ of the plot. This became the local-optimal network, denoted as *G*_0_.

### Iteratively refining the local-optimal network

In this section, we present an algorithm for refining an initial local-optimal network *G*_0_ to find a better local-optimal, or ideally, a global-optimal network *G*_*F*_ . The steps are outlined as follows:

1. Create a list of potential edges not present in the current network *G*. The number of copies *C*_*e*_ of an edge (*e*) in the list (duplicates) is calculated based on equation **6**.

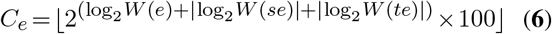

Here, *W* (*e*) represents the PPI score of edge *e, W* (*se*) denotes the fold change of the starting node of edge *e*, and *W* (*te*) signifies the fold change of the ending node of edge *e*.
2. Randomly add 0.1% of the edges from the list to *G*. Based on the number of copies obtained from equation **6**, edges with more copies have a higher probability of being added to the current network *G*.
3. Find all the significant shortest paths between each source and target pair using equations **1, 2**, and **3** (*K* = 5 and *p*(*s*) = 0.02 by default).
4. Retain the top N (5 by default) paths between each source and target pair (Equation **4**), while keeping at least *l* (8 by default) paths between each pair.
5. Determine the next local-optimal network *G*_*U*_ . Subsequently, compare *G* and *G*_*U*_ based on their graph scores (Equation **5**).
6. If *S*(*G*_*U*_) *>S*(*G*), update *G* with *G*_*U*_ and calculate the % improvement using equation **7**. Otherwise, retain *G*.

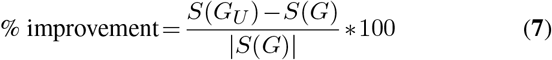
7. Repeat steps 2-6 until the average % improvement falls below 5% or there are *t* (5 by default) continuous instances of no improvement.
8. Acquire the final network *G*_*F*_ and calculate *S*(*G*_*F*_) using equation **5**.

Supplementary Figure S3 illustrates an example of the iterative refinement process, depicting the percent improvement over iterations.

### Scoring and ranking all the nodes in the network

To assess the potential consequences of each node’s removal on the globally optimized network and to subsequently identify potential intervention targets, we established a procedure to derive a ranked list of nodes. This process was initiated by the creation of a perturbed graph, denoted as *Q. Q* was generated by systematically excluding each node *n* from the final network *G*_*F*_ . Subsequently, the score *S*(*Q*) for this modified network was calculated using equation **5**. Following this, we defined the importance of each node based on the degree of reduction in the network score subsequent to its exclusion, denoted as *S*(*Q*) − *S*(*G*_*F*_). This process was repeated for all nodes in the network, consequently generating a rank order based on the magnitude of score reduction caused by each node’s removal. In this context, a node that substantially affects the network score will result in a more negative importance score (*S*(*Q*) − *S*(*G*_*F*_)), thus securing a higher rank.

The culmination of this process is an importance score, which serves as an efficacious tool for identifying and ranking key nodes (genes) within the input dataset and elucidating the implications of their inhibition or removal on the network. It is crucial to highlight that the nodes within the network *G*_*F*_ may not necessarily align with DE genes, indicating that the highest-ranked nodes (such as the top 100) might not coincide with DE genes. Therefore, our DENetwork approach augments traditional DESeq2 analyses by facilitating the identification of pivotal genes that may not be differentially expressed.

### Evaluating model performance

In assessing the performance of DENetwork, we selected the top 100 genes with the most negative importance scores in *G*_*F*_ . All top 100 genes as well as the non-DE genes within the top 100 were considered for Gene Ontology (GO) biological process (30) and Reactome pathway (31) analyses. The intersection of disease-related processes and pathways between these two sets revealed that conventional DESeq2 analyses may inadvertently neglect non-DE genes that hold relevance to disease conditions.

To illustrate the efficacy of DENetwork more tangibly, we implemented a simulation with the IAV dataset, wherein we artificially introduced receptors. This procedure was designed to demonstrate that DENetwork has the capability to accurately discern changes in genuine signal transduction occurring within the signaling pathways linking receptors, TFs, and genes. To explore the model’s robustness against noisy data, we introduced Gaussian and dropout noise to the IAV dataset. This procedure underscored DENetwork’s resilience and reliability even amidst data disruptions.

Lastly, we compared DENetwork against a well-established differential gene expression analysis method, DESeq2. This juxtaposition emphasized the pivotal role that non-DE genes play within signaling pathways, an aspect which DESeq2 may not entirely consider, thereby demonstrating the potential advantages of our approach. It is noteworthy that our benchmarking was solely against DESeq2. This choice was based on our objective to illustrate how DENetwork offers a more nuanced understanding of the transcriptomic differences between conditions, surpassing mere differential gene analysis, as typically performed by DESeq2. Further, there are limited tools available that delve into biological differences between conditions beyond differential gene expression, thereby restricting the scope for broader method comparisons.

### HuR siRNA knockdown experiments

#### Derivation and Culture of Primary Human Lung Fibroblasts

Human lung fibroblasts were derived lung tissue obtained from the University of Michigan Lung Biorepository through donor lungs provided by Gift of Life, Michigan (32) and cultured in Gibco™ Minimum Essential Media (MEM) (Thermo Fisher Scientific, USA) containing 10% fetal bovine serum (FBS; Hyclone Laboratories, Logan, UT) supplemented with gentamycin (WISENT Inc, Canada), Antibiotic-Antimycotic (WISENT Inc, Canada) and glutamax (Thermo Fisher Scientific, USA). Cells were maintained at 37°C, incubated in humidified 5% CO2-95% air and were between passages 7 to 10.

#### HuR siRNA

Fibroblasts were seeded into T25 cell culture flasks containing 4 ml of 10% FBS/MEM and allowed to grow overnight for 24 hours. Transfections were performed with either 60 nM of HuR small interfering RNA (siHuR) or control (scrambled) siRNA (siCtrl; Santa Cruz, CA). siRNA-transfected cells were incubated for an additional 24 hours and harvested.

#### RNA Sequencing

Total RNA was extracted from human lung fibroblasts using the Trizol (Bio-Rad) as per the manufacturer’s protocol. Quantification of RNA was done using Qubit (Thermo Scientific) and its quality was assessed using the 2100 Bioanalyzer (Agilent Technologies). Transcriptome libraries were generated using the KAPA RNA HyperPrep (Roche) with poly-A selection (Thermo Scientific). The starting material was 250 ng of total RNA. The library preparation steps included poly-A selection, RNA fragmentation, first and second strand cDNA synthesis, A-tailing, adapter ligation, and library amplification. Adapters used were xGen Dual Index UMI Adapters (IDT) with unique barcodes. Library quality was assessed using the 2100 Bioanalyzer, and quantification was done using Qubit and qPCR. Equimolar amounts of libraries were pooled and sequenced on the Illumina NextSeq500 platform, obtaining approximately 20M single-end 75bp reads per sample. RNA-Seq was conducted at the Institute for Research in Immunology and Cancer of the University of Montreal. Data quality was assessed with FastQC and MultiQC. Alignment was performed using STAR 2.7.8a, and non-uniquely mapped reads were discarded with Samtools 1.12. Picard Tools 2.23.3 were used for alignment quality assessment and PCR duplication rate analysis. Gene expression was quantified using featureCounts from Subread 2.0.1, and DESeq2 was employed for differential expression analysis.

### Data and Software availability

The DENetwork software, along with a comprehensive user manual and illustrative examples, can be accessed freely on GitHub at https://github.com/mcgilldinglab/DENetwork. In addition, all processed RNA-seq datasets utilized in this study are made available for reference and further exploration.

## RESULTS

### DENetwork Unveils Key Genes, DE and non-DE, Orchestrating an Influenza (IAV) Infection

To shed light on the DE gene network linked to influenza, we employed our DENetwork method on an RNA-seq dataset segregated into healthy controls and influenza-infected individuals. The dataset, after rigorous filtering and preprocessing, comprised 26,737 genes, setting up the gene universe for subsequent network analysis. H1N1 receptors, derived from a curated list (25), were earmarked as source nodes, while upregulated DE genes identified by DESeq2 served as the target nodes.

The constructed DE gene network, illustrated in Figure 2(A), contains 89 nodes (not counting 11 nodes disconnected from other top-ranked nodes). These nodes consist of 31 source nodes (receptors), 37 target nodes (upregulated DE genes), and 21 intermediate signaling proteins, each uniquely color-coded. 4 (TRIM25, DDX58, EIF2AK2, CD74) upregulated DE genes that are also receptors were designated as source nodes. Noteworthy among the larger-sized nodes, indicative of their higher ranks, are UBD (rank 1), STAT1 (rank 2), ISG15 (rank 3), TRIM25 (rank 4), and DDX58 (rank 5), all of which have previously been associated with the influenza disease (33, 34, 35, 36, 37, 38, 39) (Figure 2A).

**Figure 1.**
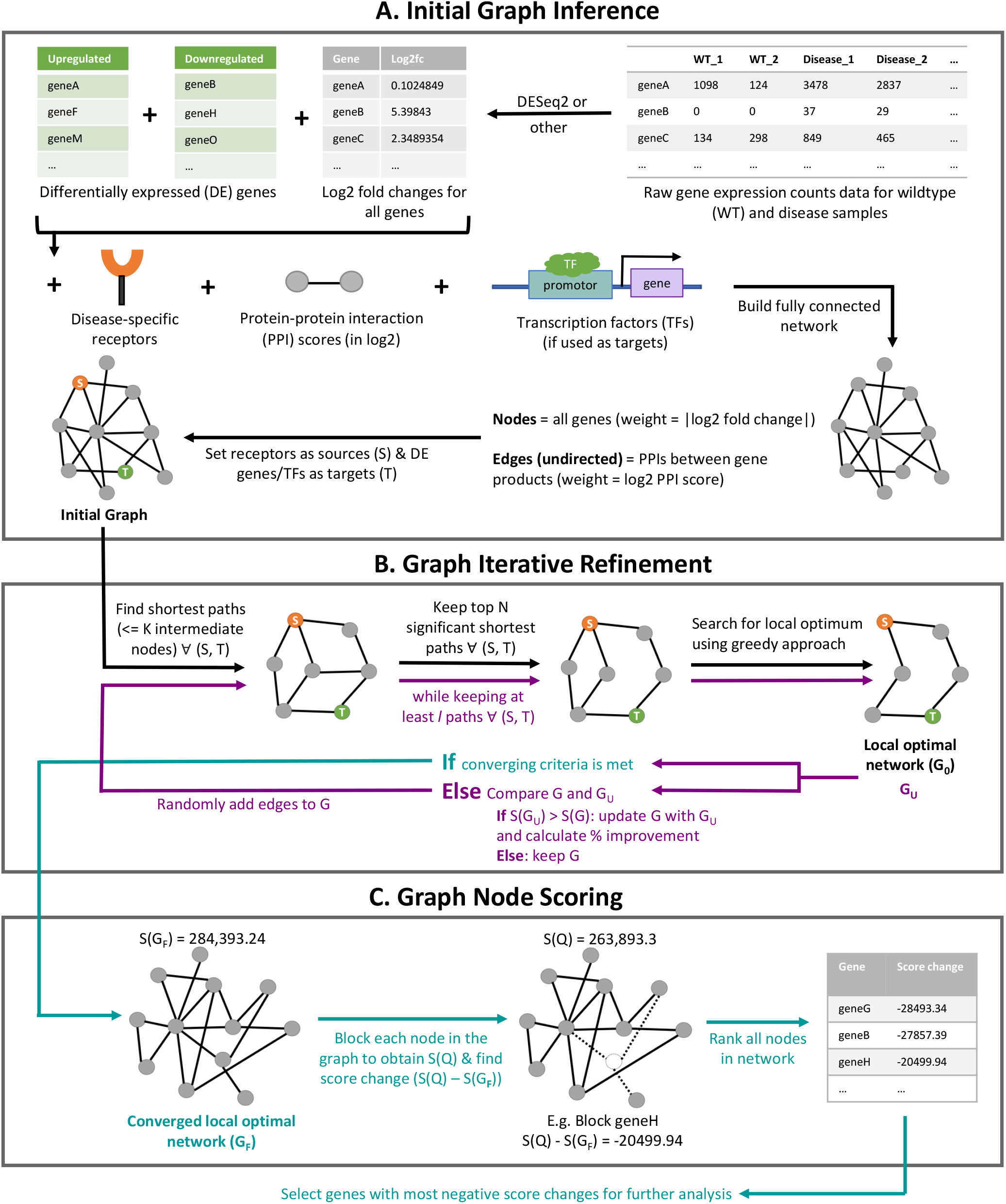
Flowchart of the DENetwork model. In **(A) Initial Graph Inference**, a fully connected network is initialized with the nodes in the graph representing the genes and the undirected edges representing the PPIs between the gene products. Disease-specific receptors are set as the source nodes (S), and DE genes or TFs are set as the target nodes (T) in the graph. In **(B) Graph Iterative Refinement**, the black arrows are followed to find the local optimal network (*G*_0_). Then, the purple arrows are followed, starting at ‘Randomly add edges to G’, to iteratively refine the network. Once the converging criteria is met, the turquoise arrows are followed to determine the converged local optimal network (*G*_*F*_). In **(C) Graph Node Scoring**, each node in the final network is blocked to find the score change between the network without the node and the original network. These score changes are used to rank the nodes, and the genes with the most negative score changes are selected for further analyses.

**Figure 2.**
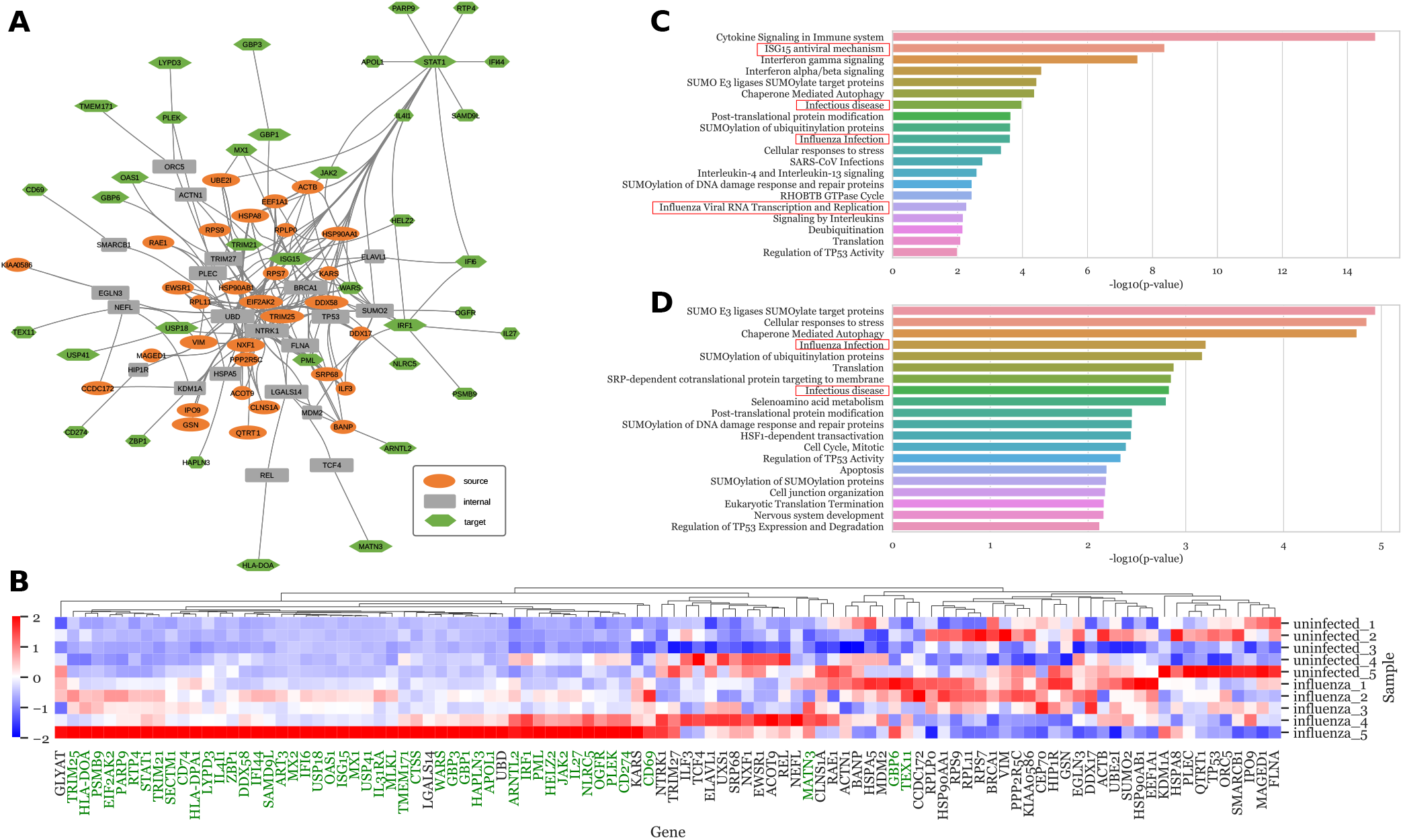
DENetwork Analysis Uncovers IAV-Associated Genes. **(A)** The graph visualizes the top 100 genes within the final network. Nodes not contributing to network connections (9 receptors/targets: MLKL, GLYAT, MX2, CEP70, ART3, IL31RA, UXS1, SECTM1, CTSS; and 2 solely interconnected nodes: HLA-DPA1, CD74) are excluded. Source nodes (receptors), internal nodes, and target nodes (upregulated DE genes) are denoted by orange ellipses, grey rectangles, and green hexagons, respectively. 4 upregulated DE genes that are also receptors were designated as source nodes. A larger node width signifies a higher network impact ranking. **(B)** A heatmap shows the z-score normalized count data of the top 100 genes across five uninfected (uninfected *) and five IAV-infected samples (infected *). Upregulated DE genes are highlighted in green. **(C)** Reactome pathway enrichment among the top 100 network genes. **(D)** Reactome pathway enrichment among the non-DE genes within the network. The top 20 Reactome pathways, ranked by FDR adjusted p-values, are depicted, with the most significant ones boxed in red. The reference set for the analysis encompasses all human genes.

Significantly, a considerable fraction of nodes (52%) within the network eludes detection by conventional differential gene analysis methodologies such as DESeq2 (colored in green in Figure 2 B). To illustrate, UBD, ranking first among nodes, and HSPA8, ranking 21st, are both non-DE genes that are implicated in IAV infections (33, 40). Furthermore, the binding of the IAV virus to HSP90AA1, a non-DE gene ranking 23rd, has been shown to facilitate viral replication (41, 42). This finding underscores the necessity of incorporating non-DE genes to gain a more comprehensive understanding of the molecular mechanisms driving IAV infections.

To deepen our understanding of the functional roles of the genes within the network, we carried out Reactome pathway enrichment analysis. The results, depicted in Figure 2C and D, reveal substantial enrichment of IAV-related Reactome pathways among the nodes of the reconstructed DE gene network. Specifically, the top 100 genes within the network participate in IAV-specific pathways such as ‘ISG15 antiviral mechanism,’ ‘Infectious disease,’ ‘Influenza Infection,’ and ‘Influenza Viral RNA Transcription and Replication’ (boxed in red in Figure 2C). Furthermore, significant enrichment of immune-related pathways such as ‘Cytokine Signaling in the Immune System’ was observed. Intriguingly, the pathway analysis of the top non-DE genes also unveiled key IAV-specific pathways, notably ‘Influenza Infection’ and ‘Infectious disease’ (boxed in red in Figure 2D). This overlap of IAV-specific pathways further accentuates the importance of non-DE genes in understanding the molecular underpinnings of IAV infections.

### DENetwork identifies critical genes driving a SARS-CoV-2 Infection

We employed DENetwork to extract a DE gene network from a SARS-CoV-2 RNA-seq dataset consisting of healthy control (neg) and diseased (pos) samples. After filtering and preprocessing, the dataset contained 27,984 genes, which constituted the gene universe for the network. The curated list of SARS-CoV-2 receptors (26) served as the source nodes, while the upregulated DE genes identified by DESeq2 served as the target nodes of the network. Figure 3A presents the reconstructed DE gene network, comprising 89 nodes (excluding 11 unconnected nodes). The network consists of 20 source nodes (receptors), 39 target nodes (upregulated DE genes), and 30 intermediate signaling proteins. 8 (LARP7, CSNK2A2, MYCBP2, PLEKHF2, MRPS27, NEK9, G3BP1, MOGS) upregulated DE genes that are also receptors were designated as source nodes. The top-ranked nodes in the network are ESR2 (rank 1), TRIM25 (rank 2), JUN (rank 3), EGFR (rank 4), and UBD (rank 5). These nodes have been previously reported to be associated with SARS-CoV-2 infections (43, 44, 45, 46, 47, 48, 49). Traditional differential gene analysis techniques often overlook source nodes and intermediate nodes that may not be differentially expressed. However, 51% of the nodes in our network are non-DE genes. Furthermore, our network analysis identifies non-DE genes that play crucial roles in SARS-CoV-2 infections (Figure 3B). For example, all the top 5 ranked nodes, despite not being DE genes, have known associations with SARS-CoV-2 infections (43, 44, 45, 46, 47, 48, 49). Furthermore, the product of NPTX1, a source node ranking 18th in the network, has been reported to interact with SARS-CoV-2 (50, 51, 52). Pathway enrichment analysis revealed significant associations of the identified nodes within the network to SARS-CoV-2-related pathways (Figure 3C and D). The pathway analysis of the top 100 genes revealed SARS-CoV-2-specific pathways such as ‘SARS-CoV-2 Infections’ and ‘Defective Intrinsic Pathway for Apoptosis’ (53, 54), as well as an immune-related pathway such as ‘Cytokine Signaling in the Immune System’. Applying this analysis to the top non-DE genes also revealed critical SARS-CoV-2-specific pathways, including ‘Defective Intrinsic Pathway for Apoptosis’, ‘Oxidative Stress Induced Senescence’, and ‘Activation of the AP-1 family of transcription factors’ (53, 54, 55, 56, 57, 58). Furthermore, TP53, a non-DE gene ranking 44th in the network, has been reported as a potential drug target for patients infected with COVID-19 (59, 60). These results highlight the importance of including non-DE genes for a comprehensive understanding of SARS-CoV-2 infections and for identification of potential therapeutic drugs.

**Figure 3.**
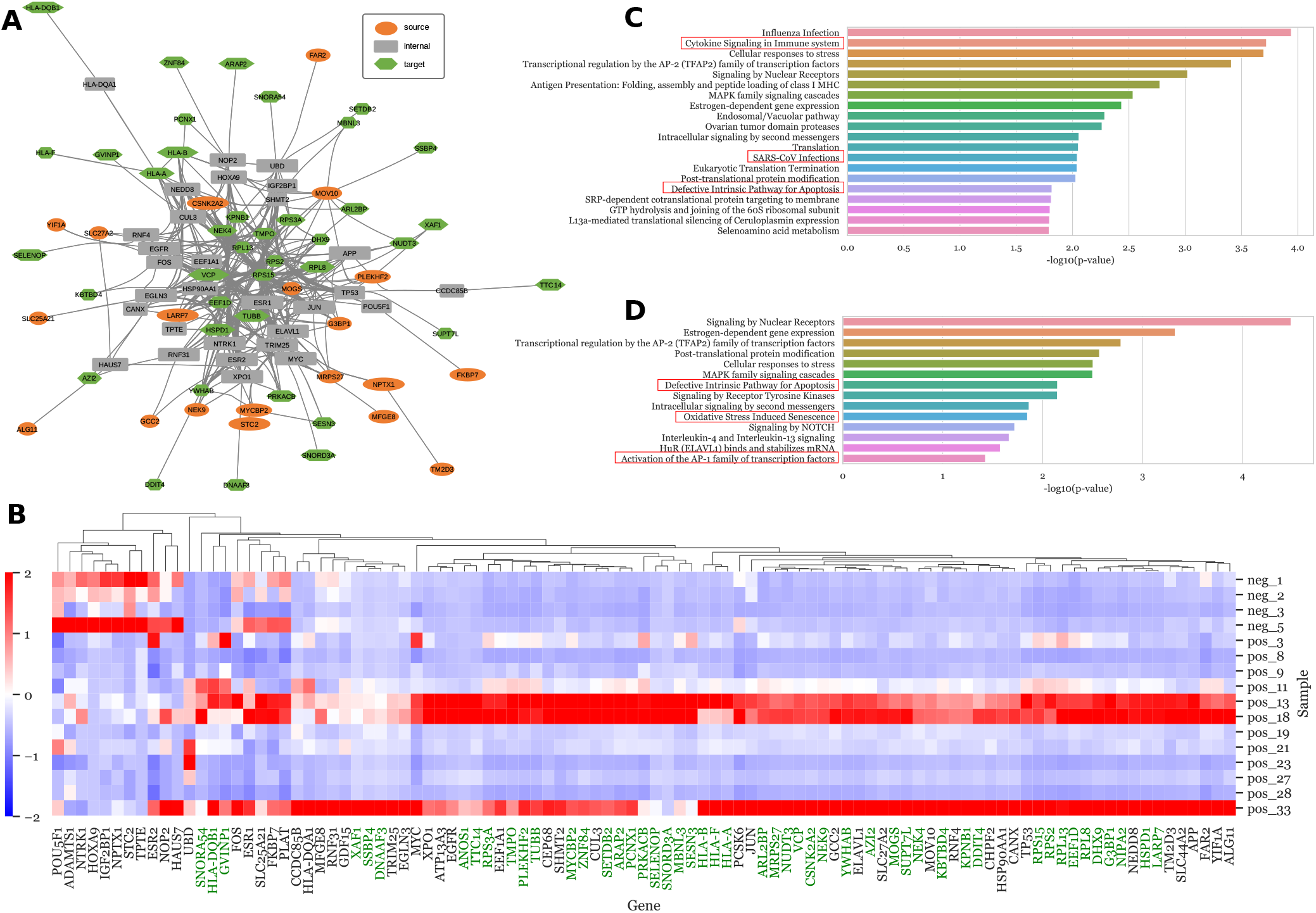
DENetwork identifies crucial DE and non-DE genes associated with COVID infection. **(A)** Network graph of the top 100 genes in the final network, with source nodes (receptors), internal nodes, and target nodes (upregulated DE genes) represented by orange ellipses, grey rectangles, and green hexagons, respectively. 8 upregulated DE genes that are also receptors were designated as source nodes. 11 source and target nodes (ATP13A3, PCSK6, ADAMTS1, SLC44A2, GDF15, CEP68, C1ORF50, ANOS1, CHPF2, PLAT, NIPA2) with no connections within the network were removed. A larger node width signifies a higher network impact ranking. **(B)** Heatmap of the normalized count data (using z-scores) for the top 100 genes across 12 positive samples (pos *) and 4 negative samples (neg *), highlighting upregulated DE genes in green. **(C)** Enriched reactome pathways for all genes in the network, using all human genes as the background. The top 20 pathways with the smallest FDR adjusted p-values are displayed, with COVID-relevant pathways highlighted in red boxes. **(D)** Enriched reactome pathways for the non-DE genes in the network. The top 20 pathways with the smallest FDR adjusted p-values are shown.

### DENetwork identifies critical genes in the macrophages of ALOX15-/-mice

To further demonstrate the effectiveness ofDENetwork, we applied it to a macrophage dataset comparing macrophages from wild-type mice and ALOX15-/-mice under two conditions: healthy (control) and diseased (22). After filtering and preprocessing, the dataset comprised 13,463 genes, which constituted the gene universe for the network. The curated receptor list for the macrophage dataset (27) was used as the source nodes, while the target nodes of the network were the upregulated DE genes identified by DESeq2. Figure 4A presents the reconstructed DE gene network, consisting of 93 nodes (excluding 7 unconnected nodes). Among these nodes, 59 are sources (receptors), 13 are targets (upregulated DE genes), and 21 are intermediate signaling proteins. 13 (ITGB7, CD36, TIMD4, ITGAE, STAB1, ITGAM, MSR1, IL1R2, C3AR1, NLRC5, LGALS3BP, NLRC3, TNFRSF13B) upregulated DE genes that are also receptors were designated as source nodes. The top-ranked nodes in the network include HSPA1A (rank 1), VCAM1 (rank 2), YES1 (rank 3), FN1 (rank 4), and CDH1 (rank 5). These nodes have been previously associated with the prolonged inflammatory response observed in ALOX15-/-mice (61, 62, 63, 64, 65, 66). Importantly, several of the top-ranked nodes, such as YES1 (rank 3), FN1 (rank 4), and EDA2R (rank 12), are non-DE genes that are associated with immune responses (63, 64, 65, 67, 68).

**Figure 4.**
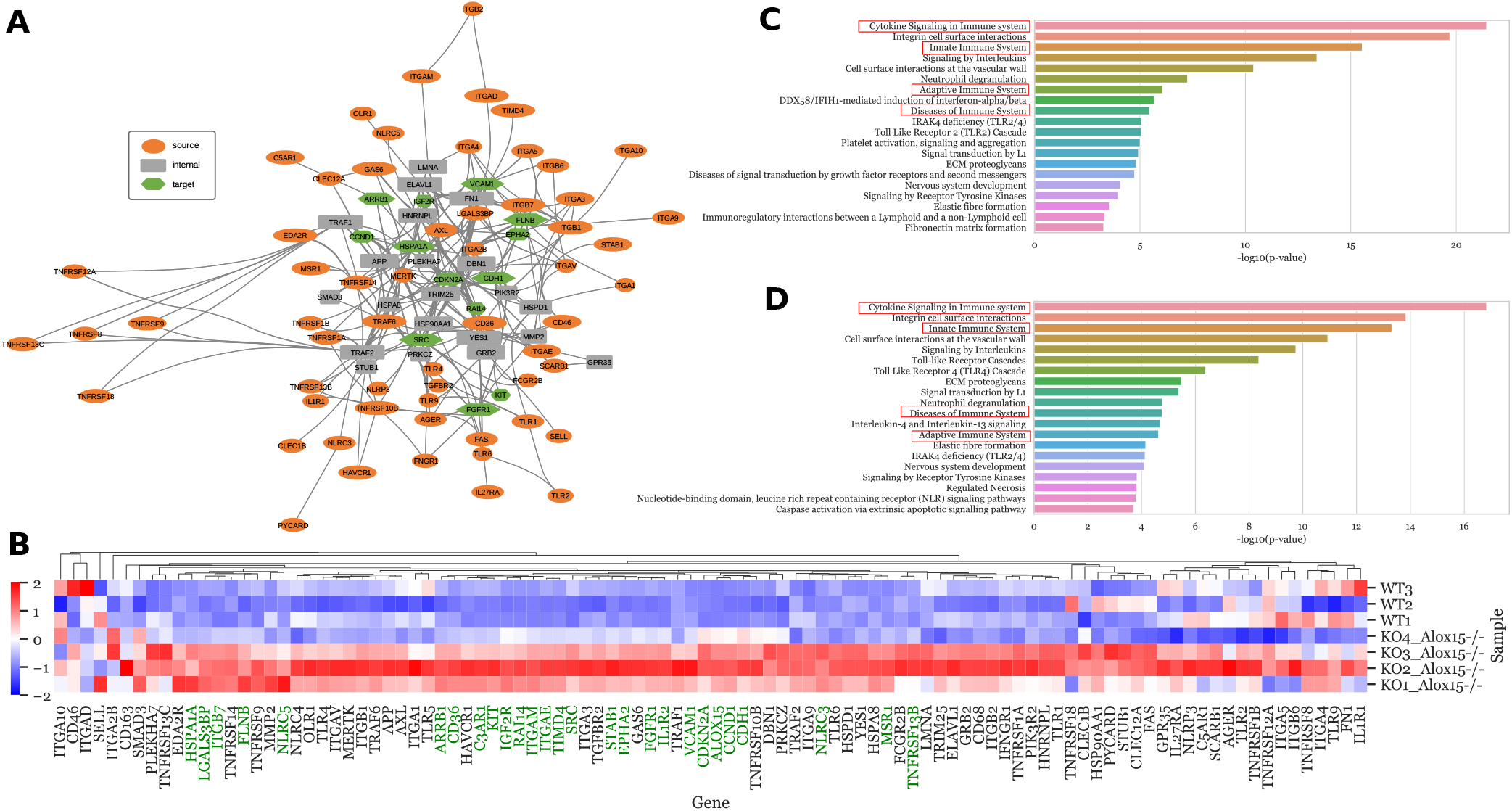
DENetwork identifies critical genes in the macrophages of ALOX15-/-mice. (**A**) Network graph of the top 100 genes in the final network. 7 (TLR5, CD68, CD163, C3AR1, ALOX15, NLRC4, IL1R2) of the top 100 genes had no connections within the network graph and were removed. Source nodes (receptors), internal nodes, and target nodes (upregulated DE genes) are represented by orange ellipses, grey rectangles, and green hexagons, respectively. 13 upregulated DE genes that are also receptors were designated as source nodes. The larger the width of the node, the higher the ranking (impact) it has in the network. (**B**) Heatmap of the normalized (using z-scores across the samples) count data of the top 100 genes across 3 wide-type samples (WT*) and 4 Alox15-/-samples (KO* Alox15-/-). Upregulated genes are colored in green.**(C)** Enriched reactome pathways for all genes in the reconstructed network. **(D)**. Enriched reactome pathways for non-DE genes in the reconstructed network. All human genes were used as the background in the analysis. The top 20 reactome pathways with the lowest FDR adjusted p-values are shown. The most relevant reactome pathways are boxed in red.

Seventy-three percent of the nodes in the reconstructed network are not identified by conventional differential gene analysis. Only 27 genes are identified as DE genes by DESeq2, and exhibit different expression patterns between the KO Alox15-/- and WT sample groups (Figure 4B). To gain further insights, we conducted reactome pathway enrichment analysis on the genes in the network. The nodes in the reconstructed DE gene network are significantly enriched in pathways associated with immune responses (Figure 4C and D). The pathway analysis of the top 100 genes reveals immune-specific pathways such as ‘Cytokine Signaling in Immune system’, ‘Innate Immune System’, ‘Adaptive Immune System’, and ‘Diseases of Immune System’ (boxed in red in Figure 4C). Notably, the pathway analysis of the top non-DE genes, which cannot be identified by DESeq2, also includes these important immune-specific pathways (boxed in red in Figure 4D). This emphasizes the significance of considering non-DE genes in ALOX15-/-mice to gain a deeper understanding of the systematic differences between the conditions.

### DENetwork identifies critical genes that drive

#### HuR-dependent pathways important for lung fibroblast function

Finally, we applied DENetwork to validate genes derived from an RNA-seq dataset experimentally obtained from human lung fibroblasts. In this dataset, the expression of human antigen R (HuR) was silenced via an siRNA technique. HuR, an RNA-binding protein encoded by the ELAVL1 gene, belongs to the Hu/embryonic lethal, abnormal vision (ELAV) family (69). HuR plays a critical role in the stabilization of target mRNA by binding to the adenylate- and uridylate-rich element (ARE) within the mRNA’s 3’ untranslated region (3’UTR) (70). This stabilization enhances and allows for the mRNA to be translated into protein. HuR is present in a variety of lung cells, including epithelial cells and fibroblasts (71, 72), and it modulates lung fibroblast differentiation and extracellular matrix (ECM) production in response to transforming growth factor-*β* (TGF-*β*) (23, 24). However, there remains a lack of knowledge regarding the extent to which HuR regulates fundamental biological processes in primary lung cells, information that could greatly assist in the identification of novel therapeutic targets for fibrotic lung diseases.

Fibroblasts are multifunctional mesenchymal cells found in various tissues, including the lung, which play vital roles in homeostasis, development, and wound healing (73). They contribute to ECM production by providing structural support, and are implicated in initiating immune and angiogenic responses. These responses are crucial in preserving tissue integrity and promoting regeneration following injury (74). Previous research has demonstrated that HuR, through its stabilization of the transcript, regulates the differentiation of fibroblasts into myofibroblasts, the primary cell type responsible for ECM production (23, 24, 75, 76, 77, 78, 79, 80).

To further explore the role of HuR in mRNA stabilization, we used siRNA to silence HuR in primary lung fibroblasts and subsequently examined the transcriptome via RNA-seq. The DENetwork analysis of the RNA-seq dataset revealed ELAVL1 as one of the top DE genes (Fig. 5A and 5B), a finding that corroborates the achieved knockdown of over 50% of ELAVL at the mRNA level in lung fibroblasts (23). The reconstructed DE gene network, comprises 100 nodes: 82 source nodes (receptors), 6 target nodes (DE genes), and 12 intermediate signaling proteins (Figure 5A). Each category of nodes is uniquely colored for clarity.

**Figure 5.**
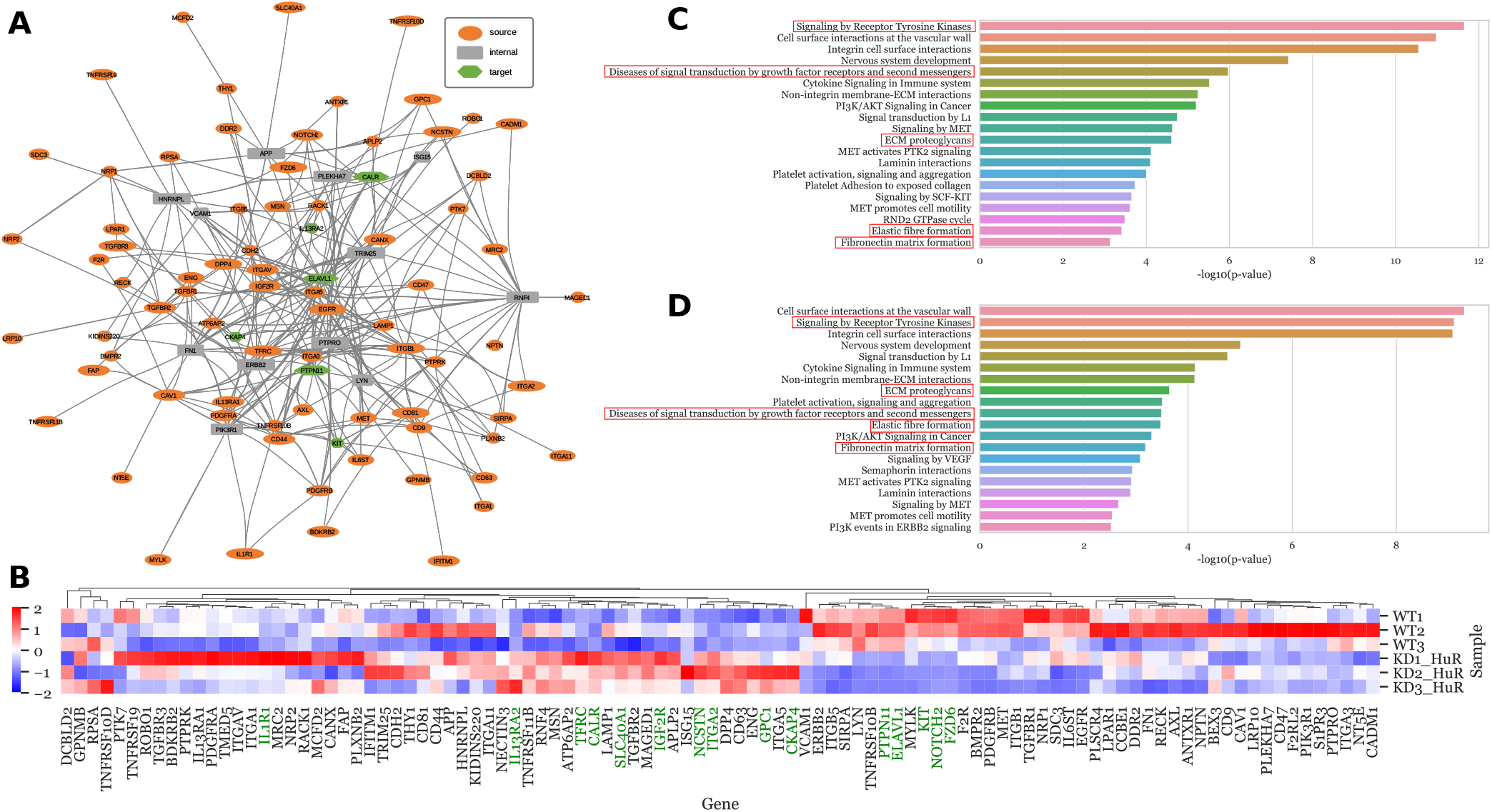
DENetwork identifies fibroblast-relevant genes and pathways controlled by HuR obtained from an experimental RNA-seq dataset. Primary human lung fibroblasts from three different subjects (1-3) underwent HuR knockdown (KD HuR) via siRNA; control lung fibroblasts received a scrambled sequence (WT). Following siRNA transfection, total RNA was isolated and RNA-seq performed as described in the Materials and Methods and the top 100 genes in the final network were selected for further analysis. **(A)** Network graph of the top 100 genes. Source nodes (receptors), internal nodes, and target nodes (all DE genes, both upregulated and downregulated) are represented by orange ellipses, grey rectangles, and green hexagons, respectively. 9 DE genes that are also receptors are used as source nodes in the network. The larger the width of the node, the higher the ranking (impact) it has in the network. **(B)** Heatmap of the normalized (using z-scores across the samples) top 100 genes in the final network across 6 samples (lung fibroblasts with control siRNA labelled WT and lung fibroblasts with HuR-knockdown labelled KD HuR). DE genes are in green, and non-DE genes identified by DENetwork are in black. **(C)** Reactome pathway annotations of all the top 100 genes are shown. **(D)** Reactome pathways identified from the top non-DE genes. The reference set for the analysis encompasses all human genes. The top 20 pathways with the lowest FDR adjusted p-values are shown. The most relevant Reactome pathways for fibroblast biology are boxed in red.

DENetwork successfully captured a significant number of non-DE genes regulated by HuR that are essential for normal fibroblast function, including TGFBR1, TGFBR2, TGFBR3, ITGA1, THY1, PDGFRB, EGFR, FN1, and PIK3R1 (81, 82, 83) (Figure 5B). Of particular interest was the capture of the ‘Signaling by Receptor Tyrosine Kinases’ pathway, identified by both DESeq2 and DENetwork (Figure 5C and D). This pathway encompasses genes pivotal to fibroblast biology as well as genes whose protein levels are controlled by HuR (e.g. FN1) (84, 85, 86, 87). Furthermore, four other pathways associated with fibroblast function (‘Diseases of signal transduction by growth factor receptors and second messengers’, ‘ECM proteoglycans’, ‘Elastic fibre formation’, and ‘Fibronectin matrix formation’) were also identified by pathway analysis of the top non-DE genes (Figure 5D). In particular, the ‘Elastic fibre formation’, and ‘Fibronectin matrix formation’ pathways also included FN1.

Our analysis also identified non-DE factors controlling FN1 production, namely MET and TGFBR2, hinting at a potential HuR-regulated mechanism governing FN1 levels. MET, a receptor tyrosine kinase that controls cell survival, migration, and invasion, has been implicated in fibrosis across various organs, underscoring its role in fibroblast-driven tissue remodeling (88, 89, 90). TGFBR2, an integral component of the TGF-*β* signaling cascade, forms a heteromeric receptor complex on the cell surface (91) that triggers fibroblast differentiation into myofibroblasts, thus enhancing ECM production. Our findings indicate that TGFBR2 expression is dependent on HuR, lending further insight into HuR’s regulatory role in the TGF-*β* signaling pathway (23).

In conclusion, the use of DENetwork to investigate HuR-dependent pathways reaffirms the existing literature on HuR regulation of fibroblast biology (23, 24, 75, 76, 77, 78, 79, 80). Moreover, it unveils new potential targets, that may not be captured by conventional differential gene analysis, for further exploration.

### DENetwork captures the true signal transduction change as demonstrated in a simulated dataset

Using the Influenza dataset, we performed a simulation to evaluate the effectiveness of our DENetwork method in identifying the implanted receptors and TFs and their influence on gene expression. The original fully connected network of the Influenza dataset consisted of 172 receptors, 296 TFs, and 15,322 genes. To simulate the dataset, we maintained the ratio of receptors, TFs, and genes from the original network. We randomly selected 5% of the receptors (8 receptors) and determined the shortest paths independently between these receptors and the 296 TFs, as well as between the 296 TFs and the 15,322 genes. Based on the total number of paths, we selected the top 5% most connected TFs (14 TFs) and the top 5% most connected genes (760 genes) with paths of length ≤ 3 (2 edges) to the TFs. Consequently, the simulated dataset consisted of 8 receptors, 14 TFs, and 760 genes, maintaining a 1/20th fraction of the original network’s components. The DE genes in the simulation comprised the downstream target genes of the implanted receptors and TFs, totaling 782 genes. In the simulation, we increased the expression of these genes by a factor of 2 (log2 fold change increase of 1.0), while keeping the expression of all other genes unchanged.

We applied the DENetwork method to the simulated RNA-seq data with perturbed expression of the 782 genes. The analysis successfully identified all inserted receptors and 13 out of the 14 TFs in the reconstructed network. Among the top 100 ranked nodes, there were 32 receptors and 32 TFs, encompassing all 8 implanted receptors and 13 out of 14 implanted TFs. The F1 scores for the recovery of the implanted receptors and TFs were 0.4 (TP = 8, FP = 32-8, FN = 0) and 0.565 (TP = 13, FP = 32-13, FN = 1), respectively. These results demonstrate the robustness of the DENetwork method in accurately identifying the essential receptors and TFs responsible for downstream changes in gene expression levels.

### DENetwork is robust against noisy data

To assess the robustness of DENetwork against noisy data, we conducted experiments on the Influenza dataset (GSE192528) by adding Gaussian noise and dropout noise to a randomly selected subset of genes. For Gaussian noise, we added noise to 5%, 10%, and 20% of the genes in the dataset. We performed 10 runs for each noise level, resulting in a total of 30 runs. In each run, we used the TFs identified using upregulated DE genes as targets for the model. To evaluate the performance, we compared the overlap percentage of the top 100 genes obtained from the noisy datasets with those from the original dataset. Our findings demonstrated that adding Gaussian noise to either 5%, 10%, or 20% of the genes preserved a median of 82-84% of the top 100 genes, illustrating the robustness of our method to noisy data (Figure 6B). While a few runs at the 5% and 10% noise levels exhibited low overlap percentages (50 for 5% and 45 and 51 for 10%), these instances were rare. Next, we examined the robustness of DENetwork against dropout noise, another common type of noise. We randomly dropped out 5%, 10%, and 20% of the genes by setting their log2 fold changes to 0. We performed 10 runs for each dropout noise level, resulting in a total of 30 runs. Using the TFs identified from upregulated DE genes as targets, we compared the overlap percentage of the top 100 genes obtained from the noisy datasets with those from the original dataset. The results showed that adding dropout noise to either 5%, 10%, or 20% of the genes maintained a median of 80-88% of the top 100 genes (Figure 6A), reaffirming the robustness of our method against noisy data.

**Figure 6.**
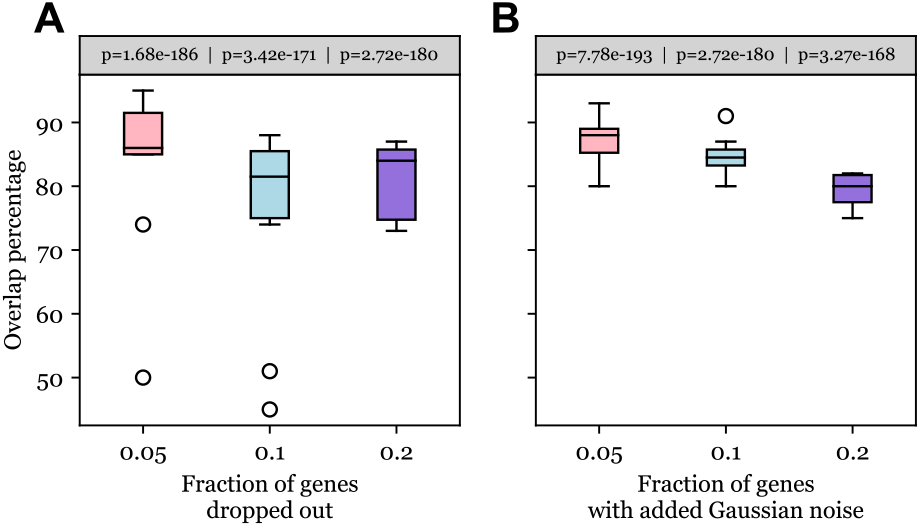
DENetwork is robust against transcriptomic noise. The boxplots represent the percent conservation of the top 100 genes within the DENetwork when subject to varying levels of simulated noise on the IAV dataset. Two prevalent types of noise found in transcriptomic measurements were simulated for this test. Panel **(A)** illustrates the resilience of the DENetwork against dropout noise, and Panel **(B)** demonstrates its robustness when exposed to Gaussian noise. The x-axis in both panels indicates the level of noise, while the y-axis signifies the robustness of the network, represented by the percent conservation of the top 100 genes under noise simulation (relative to the results without simulated noise). Noise was simulated within 5% (pink), 10% (blue), and 20% (purple) of the genes. Statistical significance of gene overlap under noise simulation, reflected by the p-value, is displayed above each corresponding box.

### DENetwork identifies important non-DE genes beyond DE genes determined by DESeq2

With an aim to clarify the role of non-DE genes in delineating different biological conditions, we strategically contrasted the results obtained from two distinct tools: DENetwork, which incorporates both DE and non-DE genes, and DESeq2, which solely identifies DE genes.

Initiating our comparative study on the IAV dataset, we identified the enriched reactome pathways associated with the top 100 genes distinguished by DENetwork and DESeq2. This initial investigation was followed by an evaluation of the statistical significance of the enriched pathways derived from both methodologies. The outcomes, illustrated in Figure 7A confirmed the superior statistical significance of the pathways enriched through DENetwork. Comparative studies on the SARS-CoV-2, macrophade, and HuR datasets also produced similar results (Supplementary Figure S4).

**Figure 7.**
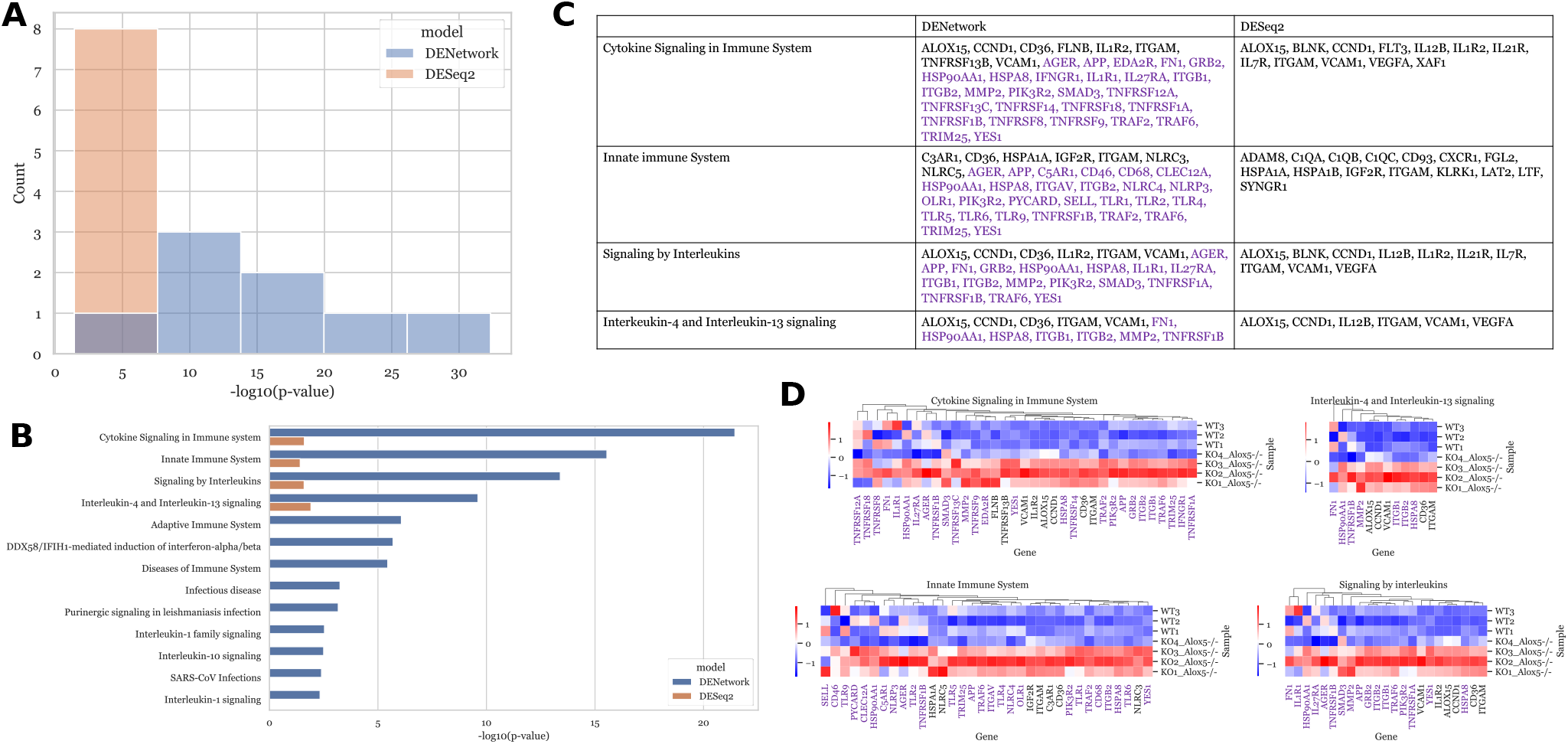
DENetwork Analysis outperforms DESeq2, the conventional differential gene analysis method. **(A)** The enriched Reactome pathways associated with the top 100 genes identified through DENetwork analysis are statistically more significant than those linked with the top 100 DE genes from DESeq2 in the sample macrophage dataset. **(B)** The enriched pathways for the top 100 genes identified by DENetwork exhibit stronger relevance to known macrophage functions such as innate immunity. **(C)** Among the top 100 genes, DENetwork identifies a greater number of genes involved in the pertinent Reactome pathways than DESeq2. All non-DE genes are colored in purple, and DE genes are colored in black. **(D)** A heatmap illustrates the gene expression of the genes involved in the shared Reactome pathways between the DENetwork and DESeq2 results. Non-DE genes, marked in purple, constitute a significant proportion, underlining the superior performance of DENetwork.

Notably, DENetwork proved its competence by revealing an extra nine significant immune-related pathways that DESeq2 failed to disclose, as shown in Figure 7B. A striking feature of the four shared top immune-related pathways between DENetwork and DESeq2 is that most of the involved genes were non-differentially expressed, colored in purple in Figures 7C and 7D. This finding underscores the potential of non-DE genes in unveiling differences among biological conditions. Given these observations, we argue that a viewpoint primarily anchored in differential gene analysis, such as the one facilitated by DESeq2 and similar tools, might inadvertently limit our comprehension of the nuanced differences between various biological samples and conditions.

## DISCUSSION

In this study, we introduce DENetwork, a graphical model aimed at advancing our comprehension of biological discrepancies between various conditions. This approach innovatively integrates gene expression measurements and established knowledge of protein-protein and protein-DNA interactome to build a signaling network that links upstream receptors to downstream DE genes identified via conventional differential gene analysis tools such as DESeq2.

The distinguishing power of DENetwork is manifested in its unique ability to pinpoint crucial non-DE genes in numerous studies of contrasting biological conditions, an achievement unattainable through traditional differential gene analysis. The superior performance of DENetwork becomes evident when compared to DESeq2, as it identifies gene sets that provide a considerably stronger statistical significance in functional enrichment analysis.

Offering a broader perspective, DENetwork challenges the prevalent inclination towards focusing solely on DE genes. Notably, our analysis shows that a substantial proportion of essential genes, including those in critically relevant pathways (e.g., COVID-relevant pathways in the COVID study), were non-differentially expressed. This observation accentuates the importance of these genes in deepening our understanding of variances across biological conditions.

In conclusion, the DENetwork framework signifies a shift in the paradigm of gene expression analysis, stressing the essential contribution of non-DE genes. This integrative approach paves the way for more precise interpretations of biological phenomena and has the potential to spark transformative breakthroughs in future research.

