## Supplementary Figures for "DENetwork: Unveiling Regulatory and Signaling Networks Behind Differentially-Expressed Genes"

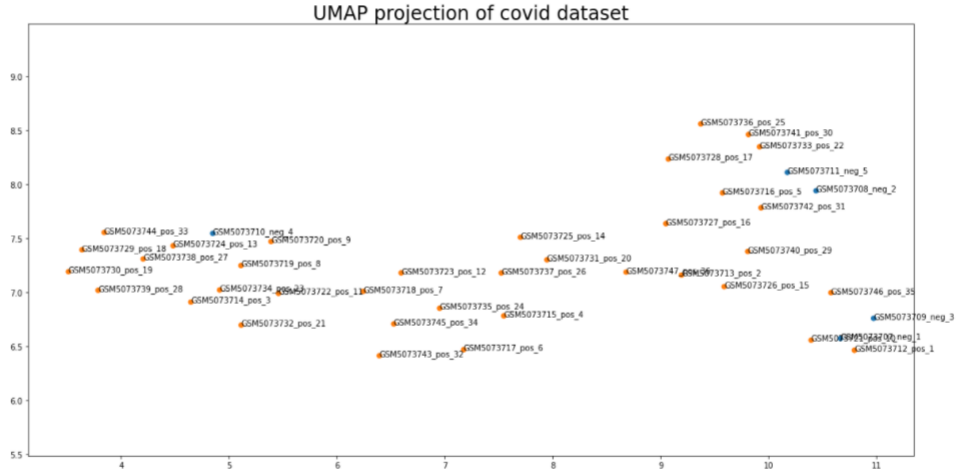


**Supplementary Figure S1.** UMAP projection of the positive and negative samples in the SARS-CoV-2 dataset. We selected the twelve positive samples on the far right and the four negative samples on the far left.

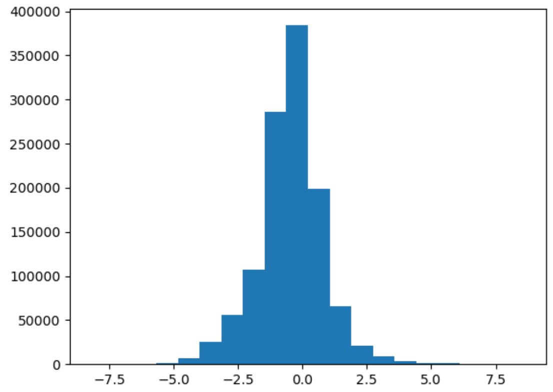


**Supplementary Figure S2.** The path score distribution (number of paths vs. path score) of the initial fully connected network of the IAV dataset. The distribution is approximately Gaussian.

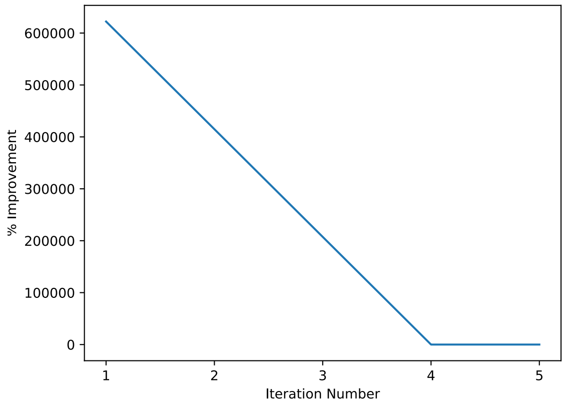


**Supplementary Figure S3.** Percent improvement vs. number of iterations plot showing the iterative refinement process of DENetwork on the IAV dataset (with TFs as target nodes). The percent improvement plateaus after the fourth iteration.

**A**

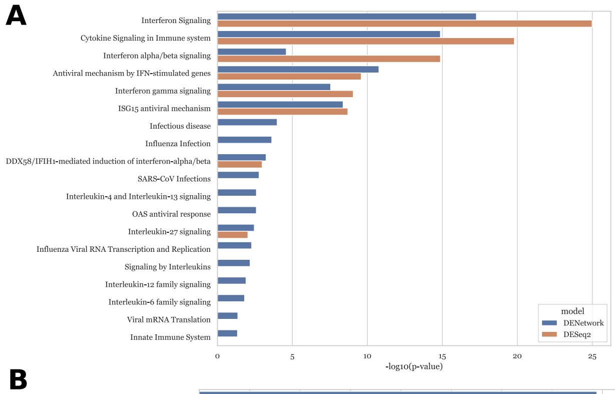


**B**

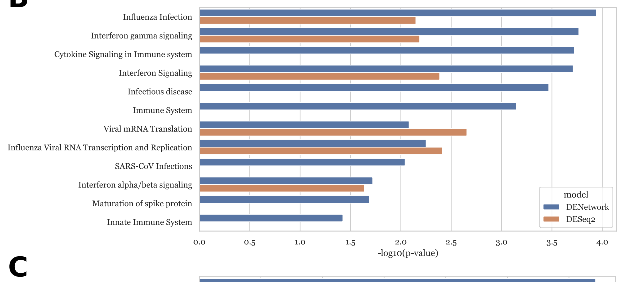


**C**

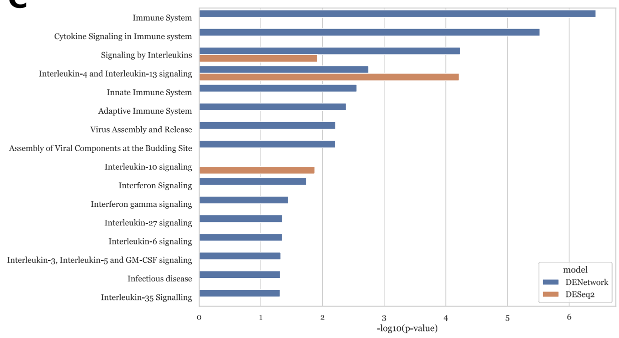


**D**

| | DENetwork | DESeq2 |
|---|---|---|
| Interferon Signaling | DDX58, EIF2AK2, GBP1, GBP3, GBP6, HLA-DPA1, IFI6, IRF1, ISG15, JAK2, MX1, MX2, OAS1, PML, STAT1, TRIM21, TRIM25, USP18, USP41, FLNA, RAE1 | AC112719, BST2, CIITA, DDX58, GBP1, GBP2, GBP3, GBP4, GBP6, HERC5, IFI35, IFI6, IRF1, IRF7, IRF8, ISG15, JAK2, MX1, MX2, OAS1, SAMHD1, STAT1, UBE2L6, USP18, USP41, XAF1 |
| Cytokine Signaling in Immune system | DDX58, EIF2AK2, GBP1, GBP3, GBP6, HLA-DPA1, IFI6, IL27, IL31RA, IRF1, ISG15, JAK2, MX1, MX2, OAS1, PML, PSMB9, STAT1, TRIM21, TRIM25, USP18, USP41, FLNA, HSP90AA1, HSPA8, RAE1, RPLP0, TP53, VIM | AC112719, BST2, CCL2, CIITA, DDX58, GBP1, GBP2, GBP3, GBP4, GBP6, HERC5, IFI35, IFI6, IL18BP, IL27, IL31RA, IRF1, IRF7, IRF8, ISG15, JAK2, LTA, MX1, MX2, OAS1, PSMB9, SAMHD1, STAT1, TNFSF13B, UBE2L6, USP18, USP41, XAF1 |
| Interferon alpha/beta signaling | IFI6, IRF1, ISG15, MX1, MX2, OAS1, USP18 | BST2, GBP2, IFI35, IFI6, IRF1, IRF7, IRF8, ISG15, MX1, MX2, OAS1, SAMHD1, USP18, XAF1 |
| Antiviral mechanism by IFN-stimulated genes | DDX58, EIF2AK2, ISG15, MX1, MX2, OAS1, STAT1, TRIM25, USP18, USP41, FLNA, RAE1 | AC112719, DDX58, HERC5, ISG15, MX1, MX2, OAS1, STAT1, UBE2L6, USP18, USP41 |
| Interferon gamma signaling | GBP1, GBP3, GBP6, HLA-DPA1, IRF1, JAK2, OAS1, PML, TRIM21, TRIM25 | CIITA, GBP1, GBP2, GBP3, GBP4, GBP6, IRF1, IRF7, IRF8, JAK2, OAS1 |
| ISG15 antiviral mechanism | DDX58, EIF2AK2, ISG15, MX1, MX2, STAT1, TRIM25, USP18, USP41, RAE1 | AC112719, DDX58, HERC5, ISG15, MX1, MX2, STAT1, UBE2L6, USP18, USP41 |
| DDX58/IFIH1-mediated induction of interferon-alpha/beta | DDX58, ISG15, NLRC5, TRIM25, HSP90AA1, HSP90AB1 | DDX58, HERC5, IFIH1, IRF7, ISG15, UBE2L6 |
| Interleukin-27 signaling | IL27, JAK2, STAT1 | IL27, JAK2, STAT1 |

**E**

| | DENetwork | DESeq2 |
|---|---|---|
| Influenza Infection | KPNB1, RPL13, RPL8, RPS15, RPS2, RPS3A, CANX, HSP90AA1, XPO1 | RPL13, RPL26, RPL8, RPS15, RPS2, RPS3A |
| Interferon gamma signaling | HLA-A, HLA-B, HLA-DQB1, HLA-F, HLA-DQA1, TRIM25 | HLA-A, HLA-B, HLA-DQB1, HLA-F, TRIM22 |
| Interferon Signaling | HLA-A, HLA-B, HLA-DQB1, HLA-F, KPNB1, XAF1, HLA-DQA1, TRIM25 | DDX58, HLA-A, HLA-B, HLA-DQB1, HLA-F, TRIM22, XAF1 |
| Viral mRNA Translation | RPL13, RPL8, RPS15, RPS2, RPS3A | RPL13, RPL26, RPL8, RPS15, RPS2, RPS3A |
| Influenza Viral RNA Transcription and Replication | RPL13, RPL8, RPS15, RPS2, RPS3A, HSP90AA1 | RPL13, RPL26, RPL8, RPS15, RPS2, RPS3A |
| Interferon alpha/beta signaling | HLA-A, HLA-B, HLA-F, XAF1 | HLA-A, HLA-B, HLA-F, XAF1 |

**F**

| | DENetwork | DESeq2 |
|---|---|---|
| Signaling by Interleukins | IL13RA2, IL1R1, PTPN11, APP, CANX, FN1, IL13RA1, IL6ST, ITGB1, LYN, MSN, PIK3R1, VCAM1 | CCND1, COL1A2, CXCL8, HGF, HMOX1, IL13RA2, IL1R1, IL6, P4HB, PTGS2 |
| Interleukin-4 and Interleukin-13 signaling | IL13RA2, FN1, IL13RA1, ITGB1, PIK3R1, VCAM1 | CCND1, COL1A2, CXCL8, HGF, HMOX1, IL13RA2, IL6, PTGS2 |

**Supplementary Figure S4.** Comparing the reactome pathway annotations of the top 100 genes determined by running DENetwork and DESeq2 on the IAV, SARS-CoV2, and experimental datasets. The top 100 genes from DENetwork are the top-ranked genes that have the largest impact on the final network. The top 100 genes from DESeq2 are the most upregulated genes in the IAV and SARS-CoV2 datasets, and are the most differentially-expressed genes in the experimental dataset. -log10(p-value) of the relevant reactome pathway annotations amongst DENetwork and DESeq2 for the (**A**) IAV, (**B**) SARS-CoV-2, and (**C**) experimental datasets. Table of the genes involved in the shared reactome pathway annotations between DENetwork and DESeq2 for the (**D**) IAV, (**E**) SARS-CoV2, and (**F**) experimental datasets. Non-DE genes are colored in purple.

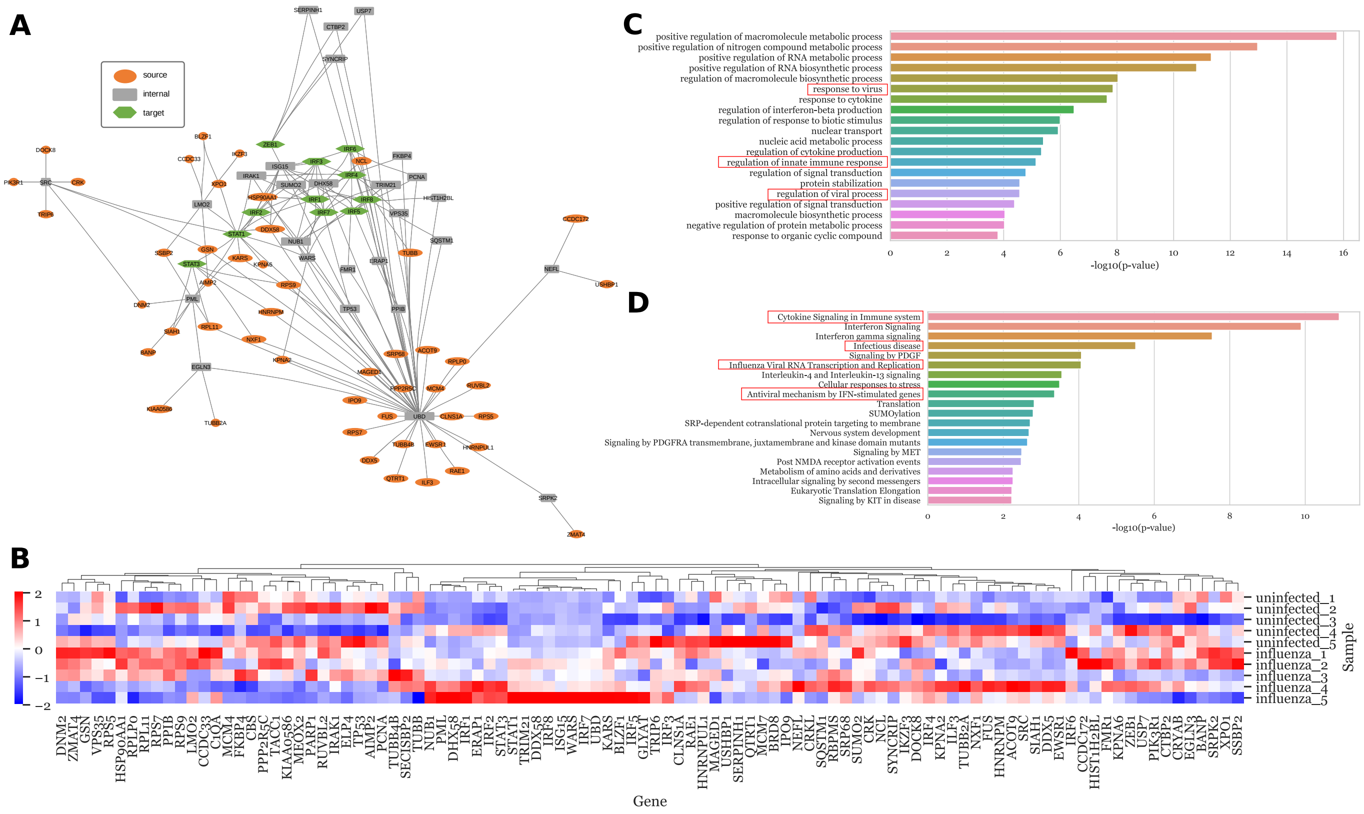


**Supplementary Figure S5.** DENetwork results on the IAV dataset with TFs (found using upregulated DE genes) as target nodes. (**A**) Network of the top 100 genes in the final graph. 13 (MCM7, CBS, ELP4, BRD8, C1QA, RBPMS, SECISBP2, PARP1, GLYAT, CRYAB, TACC1, MEOX2, CRKL) of the top 100 genes had no connections within the network and were removed. Source nodes (receptors), internal nodes, and target nodes (TFs) are represented by orange ellipses, grey rectangles, and green hexagons, respectively. Any TFs that can also be receptors are listed as receptors. A larger node width signifies a higher network impact ranking. (**B**) Heatmap of the normalized (using z-scores across the samples) counts data of the top 100 genes across 5 uninfected (uninfected_*) and 5 infected IAV samples (infected_*). (**C**) GO biological process level 3 annotations of the top 100 genes. (**D**) Reactome pathways level 3 annotations of the top 100 genes. All human genes were used as the background in the analysis. The top 20 GO biological processes and reactome pathways with the lowest FDR adjusted p-values are shown. The most relevant processes and pathways are boxed in red.

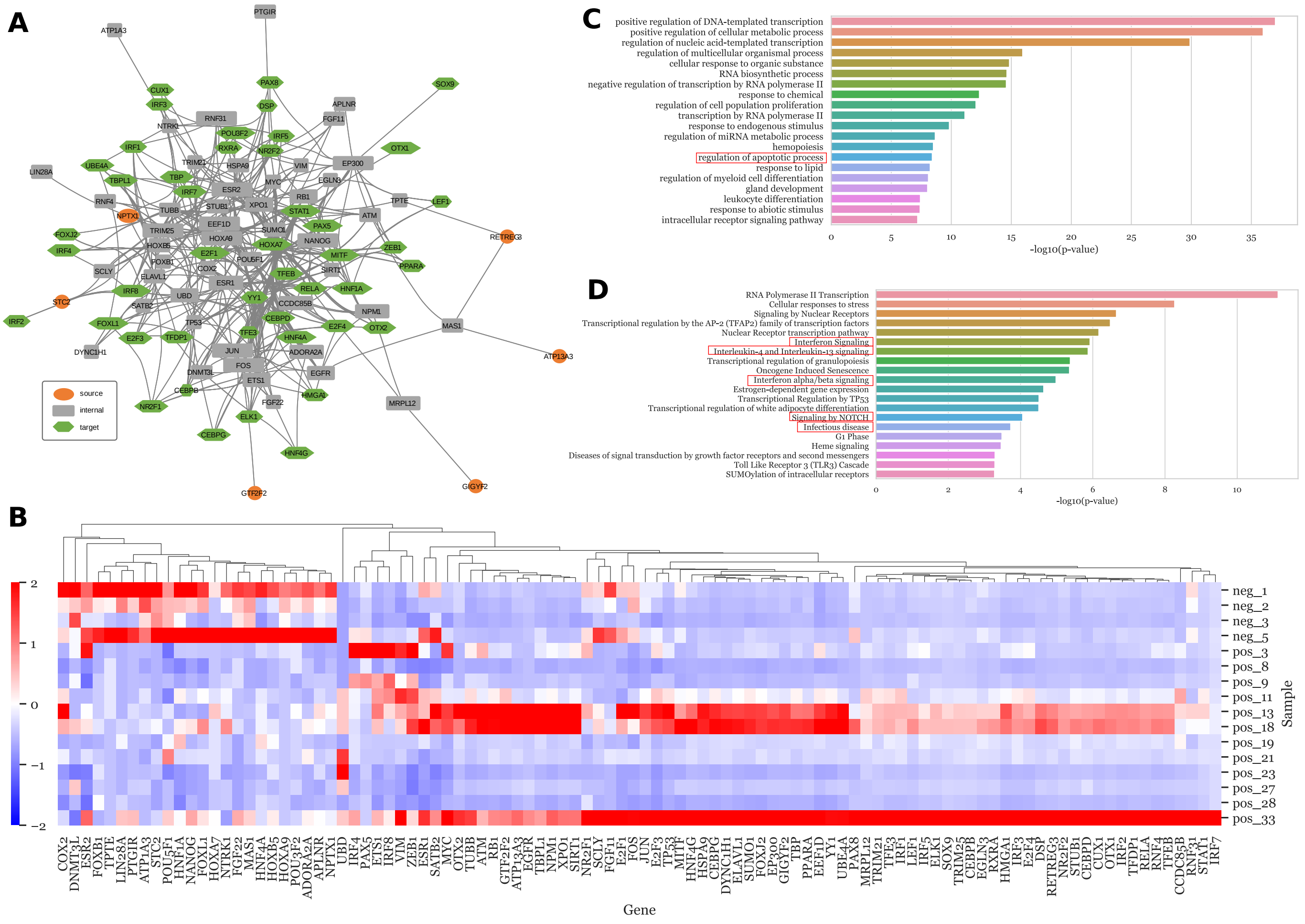


**Supplementary Figure S6.** DENetwork results on the SARS-CoV-2 dataset with TFs (found using upregulated DE genes) as target nodes. (**A**) Network of the top 100 genes in the final graph. Source nodes, internal nodes, and target nodes are represented by orange ellipses, grey rectangles, and green hexagons, respectively. Any TFs that can also be receptors are listed as receptors. A larger node width signifies a higher network impact ranking. (**B**) Heatmap of the normalized (using z-scores across the samples) counts data of the top 100 genes in the global optimal graph across 12 positive samples (pos_*) and 4 negative samples (neg_*) (**C**) GO biological process level 3 annotations of the top 100 genes. (**D**) Reactome pathways level 3 annotations of the top 100 genes. All human genes were used as the background in the analysis. The top 20 GO biological processes and reactome pathways with the lowest FDR adjusted p-values are shown. The most relevant processes and pathways are boxed in red.
